# Anti-allergic and anti-inflammatory activities of black cumin extracts in *in vitro* and *in vivo* model systems

**DOI:** 10.1101/2021.06.05.447226

**Authors:** Nazma Shaheen, Afiatul Azam, Amlan Ganguly, Saeed Anwar, Md. Sorwer Alam Parvez, Ujwal Punyamurtula, Md. Kamrul Hasan

## Abstract

Black cumin (*Nigella sativa*) is a widely used ingredient of traditional medicine for its broad-spectrum pharmacological actions, including anti-allergic, bronchial asthma, and anti-inflammatory properties. We sought to evaluate BC extracts’ efficacy for their anti-allergic and anti-inflammatory properties using a comprehensive *in vitro, in vivo*, and *silico* experimental setup. To investigate whether BC extract has anti-inflammatory, anti-allergic, and analgesic therapeutic potentials *in vitro* and *in vivo*. The activity of BC was assessed through anti-allergic activity on rat basophilic leukemia-2H3 cell line, anti-inflammatory activity on J774.1A cell line, anti-inflammatory activity by carrageenan-induced rat paw edema, analgesic activity by acetic acid-induced writhing test, and ingenuity analysis of the BC extracts in inflammation control. BC exerted potent anti-allergic activity by inhibiting antigen-induced degranulation. An anti-inflammatory effect is shown by inhibiting TNF-α production. The acetic acid-induced writhing test shown a dose-dependent reduction of writhing number following BC administration. Rat paw edema test shown the dose-dependent reduction of paw edema volume following BC administration. Ingenuity Pathway Analysis (IPA) suggested BC extracts containing ferulic acid, p-coumaric acid, kaempferol, and quercetin can inhibit inflammation. This study suggests that bioactive compounds in BC extract act as an anti-allergic and anti-inflammatory agent by regulating several downstream and upstream inflammation pathways.

## Introduction

Inflammation is a fundamental part of the body’s physiological defense mechanisms against pathogenic infections and toxic substances [1]. It is involved in the body’s response to both the initial cause and the consequences of an injury. Often, however, inflammations can be triggered inappropriately, leading to tissue destruction, which in turn can result in a range of inflammatory disorders, including rheumatoid and gouty arthritis, psoriasis, and Crohn’s disease [2]. Many studies suggest that a persistent inflammatory condition can be pervasive and develop into more clinically severe afflictions, including cardiovascular disease and cancer, often with fatal outcomes [3] [2] [4].

Current management strategies for inflammatory diseases include medications, relaxation, exercise, and surgery to correct joint damage. The medications used to treat inflammatory diseases, such as non-steroidal anti-inflammatory drugs, corticosteroids, cyclophosphamide, hydroxy-chloroquine, and biologic drugs, possibly minimize disease progression by lowering joint pain, swelling, and the inflammation itself [5] [6] [7] [8]. The type of management strategy is contingent on many factors, including the patient’s age, medical background and comorbidity, immunity status, and the severity of the symptoms of inflammatory disease [5] [8]. However, the efficacy of these management strategies can be questionable, and even if the efficacy is satisfactory, many of these strategies are often not suitable for all patients because of associated side effects [9]. Both steroidal and non-steroidal inflammatory drugs are associated with a high range of adverse effects in the long-term [10] [11]. There is thus a constant push toward the development of an effective and curative therapeutic strategy to treat inflammatory diseases.

Medicinal plants are an essential source of bioactive compounds with potential therapeutic efficacy [12]. Pharmacological investigations of medicinal plants can yield primers for the effective treatment of inflammatory diseases [12]. Black Cumin (BC, *Nigella sativa* L.) is a well-known medicinal plant used extensively in Unani, Ayurvedic, and Siddhi medicine for centuries [13]. This herb, endemic to South Asian and Mediterranean countries, is rich in bioactive compounds, including tocopherols, alkaloids, saponins, and vitamins A and C, all of which contribute to its biological functionality [14] [15] [16]. Overwhelming evidence indicates the presence of bioactive ingredients in BC that can counteract the underlying pathophysiology of many diseases, including cancers, inflammatory conditions, cardiovascular defects, and autoimmune disorders [14] [17] [18]. Previous studies have highlighted the probable anti-inflammatory and anti-analgesic activities of BC [19] [20]. However, only a limited number of studies have explored BC’s anti-inflammatory effects on subacute and chronic models of inflammation [20].

As a result, we sought to investigate the dose-response effects of the anti-inflammatory activity of BC in rats and mice with *in vitro* and *in vivo* anti-inflammatory, anti-allergic and anti-analgesic activities. Using these findings, the present study has taken an attempt to develop a probable mechanism of action for *N. sativa* through Ingenuity pathway analysis.

## Materials and Methods

### Collection and preparation of black cumin samples

Trained food sample collectors collected the samples from New market Kancha Bazar, Dhaka, Bangladesh and then critical checking of BC seed samples by the expert faculty member of the Department of Botanist, University of Dhaka. Preparation and processing of the collected samples were done using standard operating procedures. Processing involved drying of the fresh samples at 25 - 30°C, grinding by an electric blender, and preserving in an airtight container until analysis.

### Extraction procedure

Multiple rounds of sequential extraction (initially by hexane/dichloromethane (1:1 v/v) (Merck, Germany; hexane-296090, dichloromethane-270997) and then by AWA (acetone/water/acetic acid 70:29.5:0.5) (Merck, Germany; acetone-650501, acetic acid-A16283) was performed in an accelerated solvent extraction equipment known as ASE 200 (DIONEX, USA, catalogue number: 055422). A detailed description of the extraction procedure has been reported elsewhere [21] [22]. While ground samples were directly mixed with Dimethyl sulfoxide (DMSO) (Sigma Aldrich, #D-2650 Poland) to get the sample extracts for assessment of anti-allergic and anti-inflammatory activities in *in vitro* cell line models, *in vivo* assessments conducted in this study mixed dried AWA extracts with DMSO to prepare the sample extracts.

### Experimental animal

Swiss Albino mice (5 - 6 weeks of age; 20 - 30 grams) and Long-Evans rats (7 - 8 weeks of age; 100 - 130 grams) were collected from the Animal Research Branch, International Centre for Diarrheal Disease Research, Bangladesh (ICDDR, B). The animal were housed in polyvinyl cages and maintained under standard laboratory conditions (temperature 25 ± 2°C) and a 12 h light - 12 h dark cycles for seven days. The animals were fed on standard laboratory animal diet formulated by ICDDR, B and water *ad libitum*. To keep the hydration rate constant, food and water were stopped 12 h before the experiments. The Ethical and Animal Care Committee of the Institute of Nutrition and Food Sciences, University of Dhaka, Bangladesh, critically reviewed and approved this study involving in vivo models. The procedures de-scribed in this study were conducted in accordance with the Bangladesh Biosafety and Biosecurity guidelines 2019 and institutional oversight performed by qualified veterinarians [23].

### Anti-allergic activity on rat basophilic leukemia-2H3 cell line

Rat basophil leukemia (RBL)-2H3 cells [NIHS (JCRB), Tokyo, Japan: Cell No.: JCRB 0023] were used to study the anti-allergic activities of BC extracts. RBL-2H3 cells were grown in minimal essential medium (Eagle) containing 15% fetal calf serum. We then inoculated 05 × 10^5^ cells in each well of a 24-well plate and incubated at 37°C in an environment of 5% carbon dioxide for affluent growth. After overnight incubation (4 × 10^6^ cells in each well of a 24-well plate), mouse monoclonal anti-DNP IgE (Sigma Aldrich #A-6661, UK) solution was added to each of the 24 wells before incubation (for two hours at 37°C).

The cells were then incubated with 10 µl of DNP labeled human serum albumin for 30 minutes. Supernatants were separated after lysis of cells with 500 µl of Triton X - 100. The cell lysate (50 µl) along with the collected supernatant was transferred to the 96-well plate and mixed with 100 µl of 0.1 M citrate buffer containing 3.3 mM para-nitrophenyl-2-acetamide-β-D-glucopyranoside (Wako, Japan). The supernatant-cell lysate mix was then kept under incubation for 25 minutes at 37°C; it was followed by an addition (100 µl) of 2 M glycine buffer of pH 10.0 which stopped the reaction in a microplate reader and measurement of the absorbance. A stop solution was added to the lower four lanes which were used as control, and then at the same time, the substrate was added to the experimental well.

Blank without antigen and another negative control were used as a positive control with antigen and 5 µM of wortmannin (Wako, Japan) solution. Blank, negative control, and positive control were prepared by adding 490 µl of modified Tyrode’s buffer and 200 folds diluted wortmannin solution.

### Anti-inflammatory activity on J774.1A cell

Anti-inflammatory potential using J774.1A cell assay was assessed using the method as reported by Herath et al. [24]. Shortly, ground BC seed samples were mixed with DMSO to prepare 4mg/ml concentration and final strength of 40 µg/ml was employed in the assay media. For the assay, a clear supernatant was obtained by centrifuging that mixture at 3000 rpm for 5 minutes. Mouse macrophages J774A.1 cell line [NIHS (JCRB) 9108] were cultured in Dulbecco’s Modified Eagle Media (DMEM) (Sigma Aldrich **#**D-5030, USA) with 10% fetal calf serum and penicillin and streptomycin (Sigma Aldrich #F-7524, UK) at a concentration of 100 U/mL. The cells were incubated at 37°C in the incubator with an atmosphere of 5% CO2, 95% air.

The cell suspension with the concentration of 5.0 ×10^5^ cell /ml (200 µl/well) was placed in 96 well culture plates and incubated overnight (37°C, 5% CO2-95% air). Overnight cultured cells washed thrice with Hank’s solution (Sigma Aldrich #6648, UK) (37°C) after removal of culture media prior to the addition of 180 µl DMEM and followed 20 µl of 10 times diluted sample extracts (final concentration, 40 µg/ml) to the each well without lipopolysaccharides (LPS). On the other hand, 160 µl DMEM with 20 µl of LPS (final concentration, 1.0 µg/ml) after the addition of the test samples and incubated further for more than four hours. Cell supernatant was then collected for the assay of the TNF-α production, assayed by using Mouse TNF-α ELISA kit (Ready-Set-Go, eBioscience, USA). In brief, 50 µl of TNF-α capture antibody (250 times diluted) was added into 96 well plates and kept in the refrigerator (4°C) overnight. The plate was washed with 300 µl of PBS-T (0.05%Tween20 in PBS) 3 times in a sera washer (Model MW-96F., Biotech, Japan) and 100 µl of assay diluents solution was added as blocking solution. After incubation for 1hour at room temperature, the plate was washed and 50 µl of cell supernatant /standard. TNF-α was added into the wells according to plate design. Sample extract-treated cell supernatant was diluted 10 times with assay diluents, whereas, the standard was added to maintain the final concentration from 0 to 1000 pg/ml and further incubated for 2hour at RT. After at least 2 hrs incubation, plates were washed again and TNF-α detection antibody (250 times diluted) was added (50 µl) into each well and further incubated for 1hour. Avidin-HRP antibody solution (250 times diluted) was added in the same volume as earlier after washing and kept 30 minutes at RT. Finally, 50 µl of TMB-HRP substrate solution was added and incubated for 15 min., a stop solution (25 µl) was added into each well to stop the reaction and the absorbance was read in the microplate reader (model 550; Bio Rad, CA, USA) at 450 nm. From the absorbance data, the inhibition effect of food extracts on TNF-α production was calculated from the standard graph. Dose-response was done for the samples which had an inhibitory effect on TNF-α production at the concentrations of 40, 10, 3, and 1 µg/ml.

### Assessment of anti-inflammatory activity by carrageenan-induced rat paw edema

Evaluation of anti-inflammatory activity was conducted *in vivo* in the Long Evans rat model. At first 200 mg of samples (i.e., dried AWA extract) were mixed with 5.0 ml of DMSO in a shaking-incubator at 130 rpm overnight. Then the mixture was centrifuged at 3000 rpm for 5.0 minutes. Then the supernatant was collected and pipetted 500 µL of aliquots to store at 25°C.

To determine anti-inflammatory activity, edema was induced by the phlogistic agent carrageenan [25]. The control group (n=3) received normal saline per os (p.o) (10 ml/kg), while group 2 received a standard drug, Diclofenac sodium (50 mg/kg, p.o). The rest of the groups were given 800 (n=3), 400 (n=3), and 200 mg/kg body weight (n=3) p.o. of extract preparations. Thirty minutes after administering sample extracts, each rat received 0.1 ml of 1% (w/v) carrageenan (injected into the sub-plantar region of the right hind paw subcutaneously) (Sigma Aldrich, # C-1013, Germany). Paw volumes of the right hind paw of each rat were measured by plethysmometer before and 1, 2, 3, and 4 hours after carrageenan injection to determine the edema volume. The hind paw volume was evaluated for anti-inflammatory activity and expressed as % inhibition of the hind paw volume, which was calculated by the following equation:

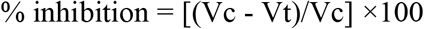

Here, Vc = average paw volume of the control group, and Vt = Average paw volume of the treated group.

### Assessment of analgesic activity by acetic acid-induced writhing test

Peripheral analgesic activity of the extracts was determined by the acetic-acid-induced writhing inhibition method in mice [26]. The control group (n=3) (group 1) received normal saline (10 ml/kg, p.o.). Group 2 received a reference drug Diclofenac sodium (50 mg/kg, p.o). The rest of the groups received 800 (n=3), 400 (n=3), and 200 mg/kg body weight (n=3) p.o. of dried AWA sample extracts. After 30 min of treatment, each mouse was administered intraperitoneally with 0.6% acetic (10 ml/kg). Later, the writhing numbers of each mouse were observed for 10 minutes. To evaluate the level of analgesia, writhing numbers of the sample treated mice were compared with the writhing numbers of the control groups. Percent-inhibition of writhing was calculated using the following equation:

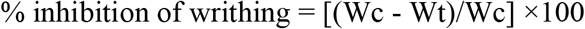

Here, Wc = Average number of the writhing of the control group, and Wt = Average number of the writhing of the treated group.

### Ingenuity analysis of the BC extracts in inflammation control

We evaluated the effects of the compounds extracted from BC using the knowledge-based path explorer feature of the Ingenuity Pathway Analysis (IPA) tool [27] [28]. We explored new ingenuity-based pathways that showed the relationship between the genes involved in the biosynthetic pathways of inflammation-related compounds (e.g., arachidonic acids, prostaglandins, and leukotrienes) and compounds in the BC extracts, which includes ferulic acid, quercetin, p-coumaric acid, and kaempferol. The shortest possible pathways were generated individually for each compound where the compounds in the BC extracts were set as the initial compound, and inflammation-related compounds were selected as the target.

## Results

### *In vitro* screening for anti-allergic effect

As compared to the negative control, BC extracts produced 46.9% (SD ±7.1) inhibition of the antigen-induced degranulation of RBL-2H3 cells. AWA extracts produced increased inhibition compared to the DMSO extracts for 5, 10, 20, and 40 µg/ml concentration. However, at higher concentrations, e.g., 40 µg/ml, both extracts produced comparable inhibition (Figure 1).

**Fig. 1.**
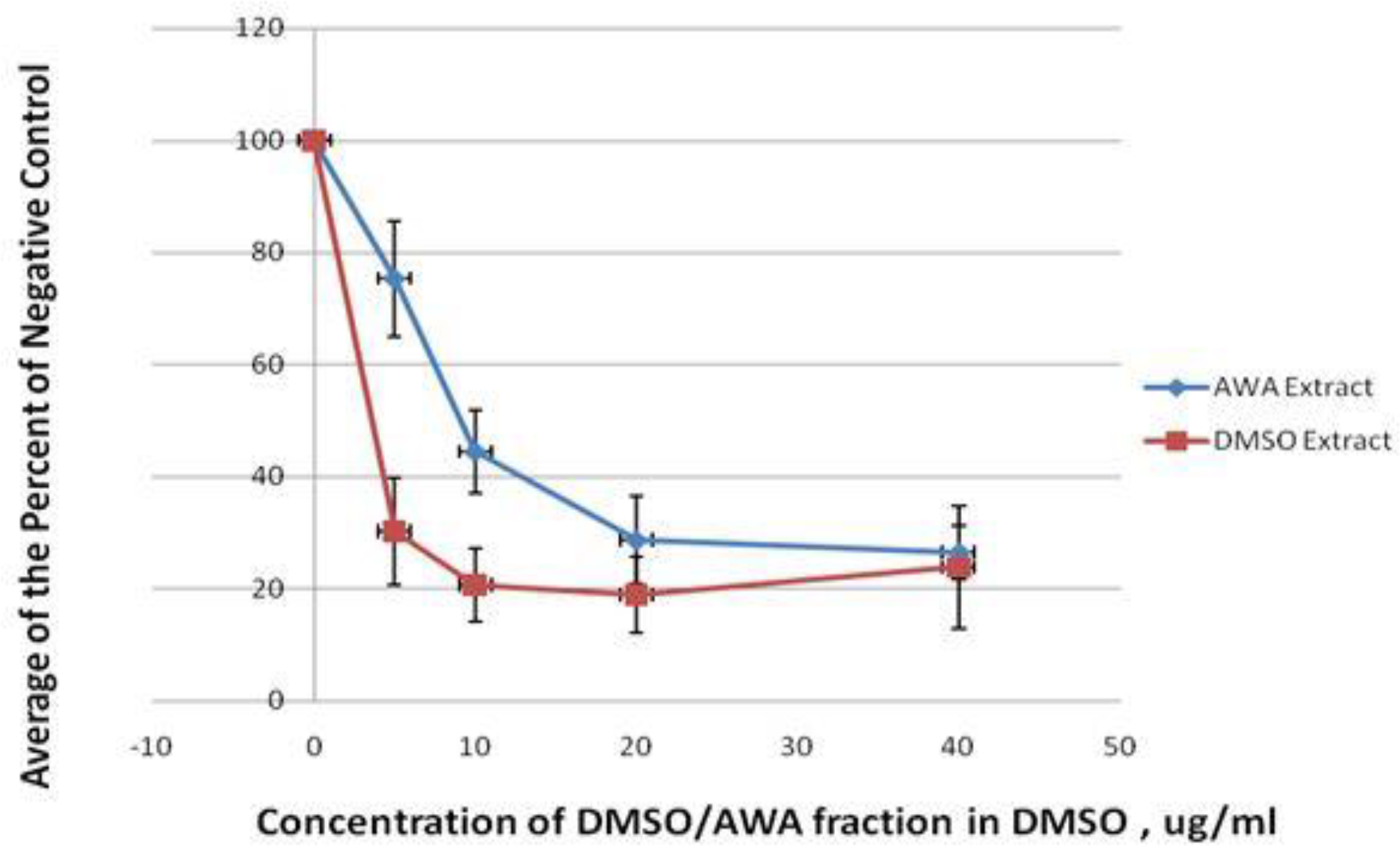
Comparison of dose-response of the antigen induced degranulation of BC extracts by DMSO (red color) and AWA (blue color) as percentages of negative control at the level of 5, 10, 20 and 40 µg/ml concentration.

### *In vitro* screening for anti-inflammatory effect

In the preliminary evaluation of anti-inflammatory effects on J774A.1 cell, 40 µg/mL of DMSO extracts of BC resulted in significant (70%) inhibition of TNF-α production. Dose-response analysis showed that 1, 3, 10, and 40 µg/mL of DMSO extracts could produce up to 70% inhibition in the TNF-α production.

### *In vivo* anti-inflammatory activity

Four hours after carrageenan injection, the maximum volume of edema in control Long Evans adult rat models was 1.18 ± 0.03 ml. Another group of rats pretreated with Diclofenac sodium (reference drug; 50 mg/kg, p.o), showed significantly reduced (p < 0.05) edema in the paw. Dose-response analysis of DMSO extract of BC produced a significant inhibition of paw edema in Long Evans adult rats in a dose-dependent manner at 200, 400, and 800 mg/kg, p.o administration of carrageenan (p<0.05) (Table 1).

**Table 1:**
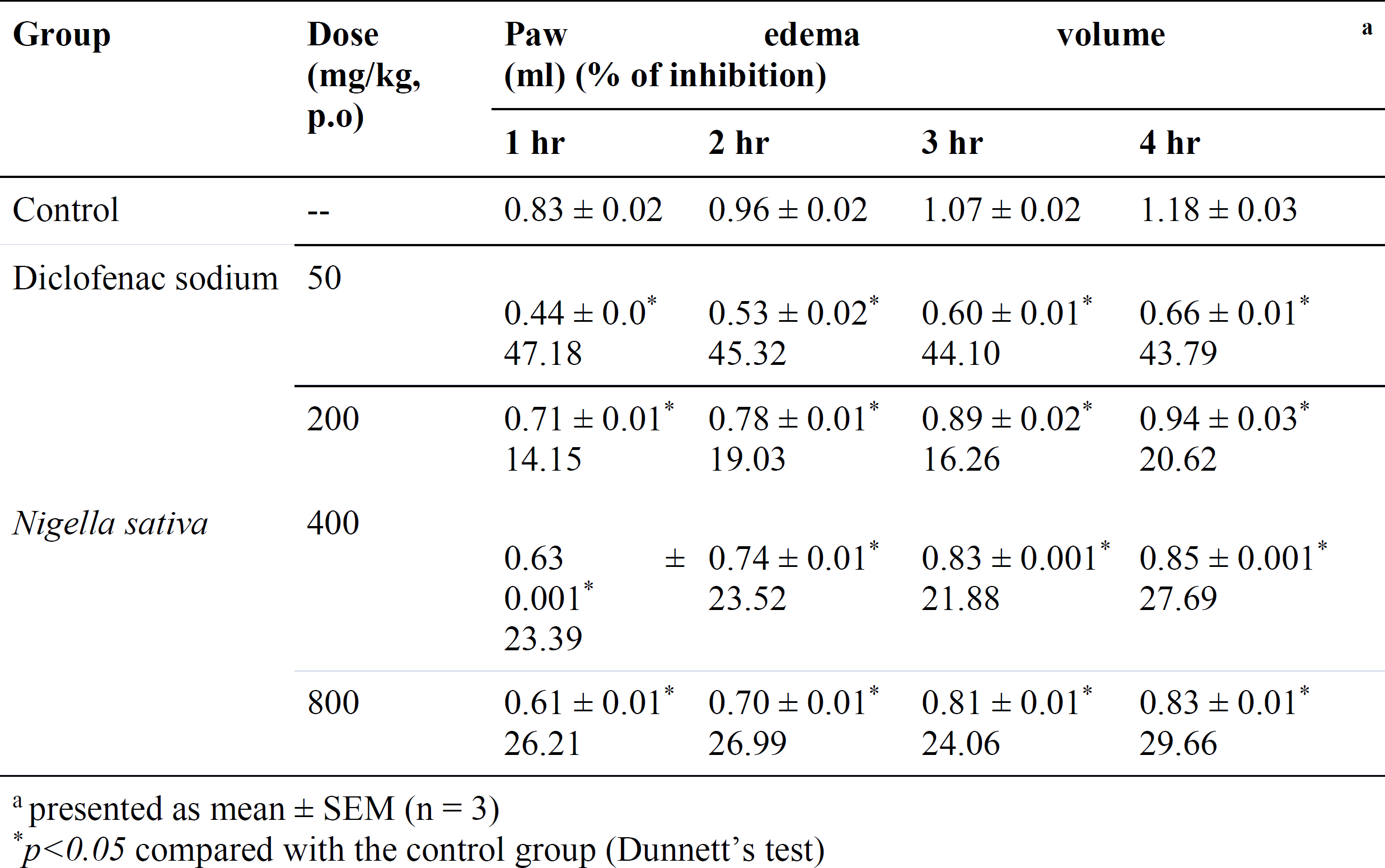
Effects Black Cumin extracts by DMSO extracts of *BC* on carrageenan-induced paw in rats

The maximum inhibition of rat paw edema by BC was noted 4 hours after administration of carrageenan at a dose of 800 mg/kg, p.o. compared to the control but less than the reference drug. Figure 2 shows the changes of percent inhibition of carrageenan-induced paw edema volume at 1, 2, 3, and 4 h at the doses of 800 mg/kg, p.o by the DMSO extract of BC when compared to the control.

**Fig. 2.**
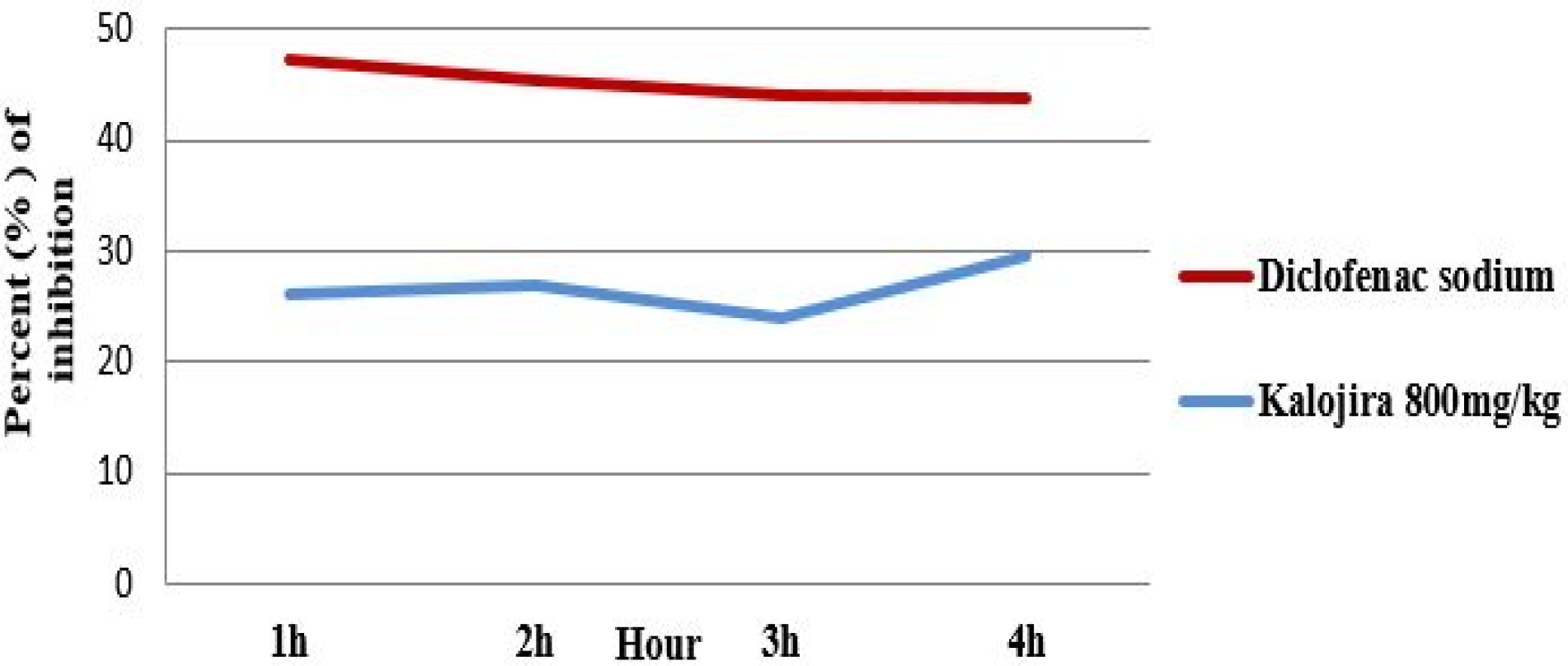
Time course in the changes of percent inhibition on carrageenan-induced paw edema volume of BC at 1, 2, 3, and 4 h at the dose of 800 mg/kg, p.o.

### *In vivo* analgesic activity

The effects of DMSO extract of BC on 0.6% acetic acid-induced writhing in Swiss albino mice are summarized in Table 2. A dose-dependent and significant (p<0.05) reduction in the number of abdominal constrictions induced by intraperitoneal administration of 0.6% acetic acid was observed with oral administration of BC, at the doses of 200, 400, and 800 mg/kg, p.o when compared to the control. Among the different doses, the DMSO extract of BC at the dose of 800 mg/kg exhibited the maximum inhibition of the number of writhing compared to that of control.

**Table 2:**
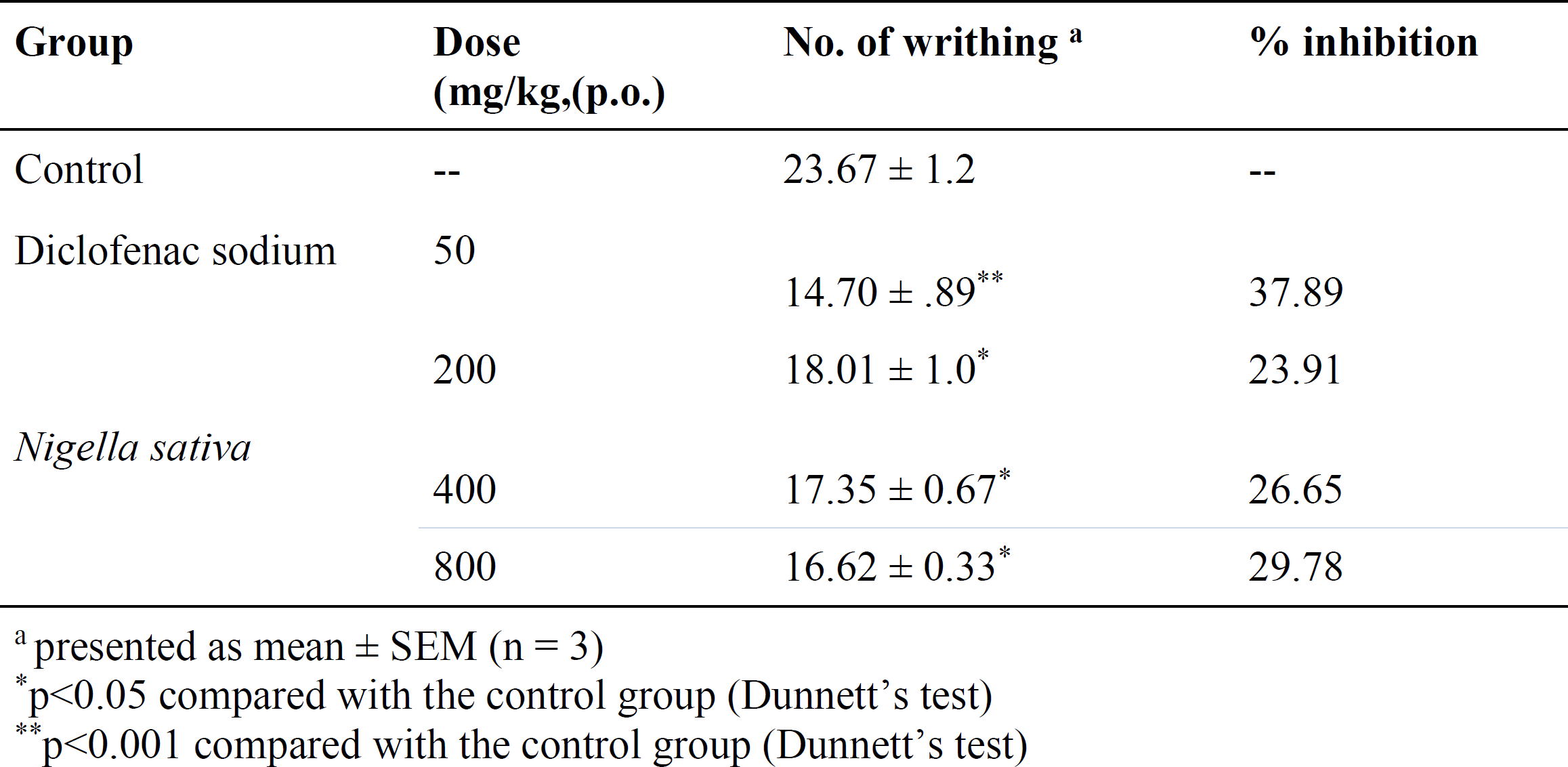
Effect of Black Cumin extracts by DMSO on acetic acid-induced (0.6%) writhing in mice

The maximum inhibition of rat paw edema by BC was noted 4 hours after administration of carrageenan at a dose of 800 mg/kg, p.o. compared to the control but less than the reference drug. Figure 2 shows the changes of percent decrease in carrageenan-induced paw edema volume at first, second, third, and fourth hours post-administration of 800 mg/kg, p.o of DMSO extract of BC when compared to the control.

### *In silico* analysis provides insights into the mechanism of how BC extracts reduce inflammation

The *in-silico* analysis of the bioactive compounds of BC illustrates a facet reactome depicting potential models of how BC extracts reduce inflammation in humans (Figure 3, S1 Table). Both prostaglandin and leukotriene synthesize from the arachidonic acid. Quercetin, p-coumaric acid, and kaempferol reduce the expression of myc, MAPK, EGFR, and TNF, all of which are involved in the upregulation of the expression of arachidonic acid. Consequently, quercetin, p-coumaric acid, and kaempferol downregulate the expression of arachidonic acids.

**Fig. 3.**
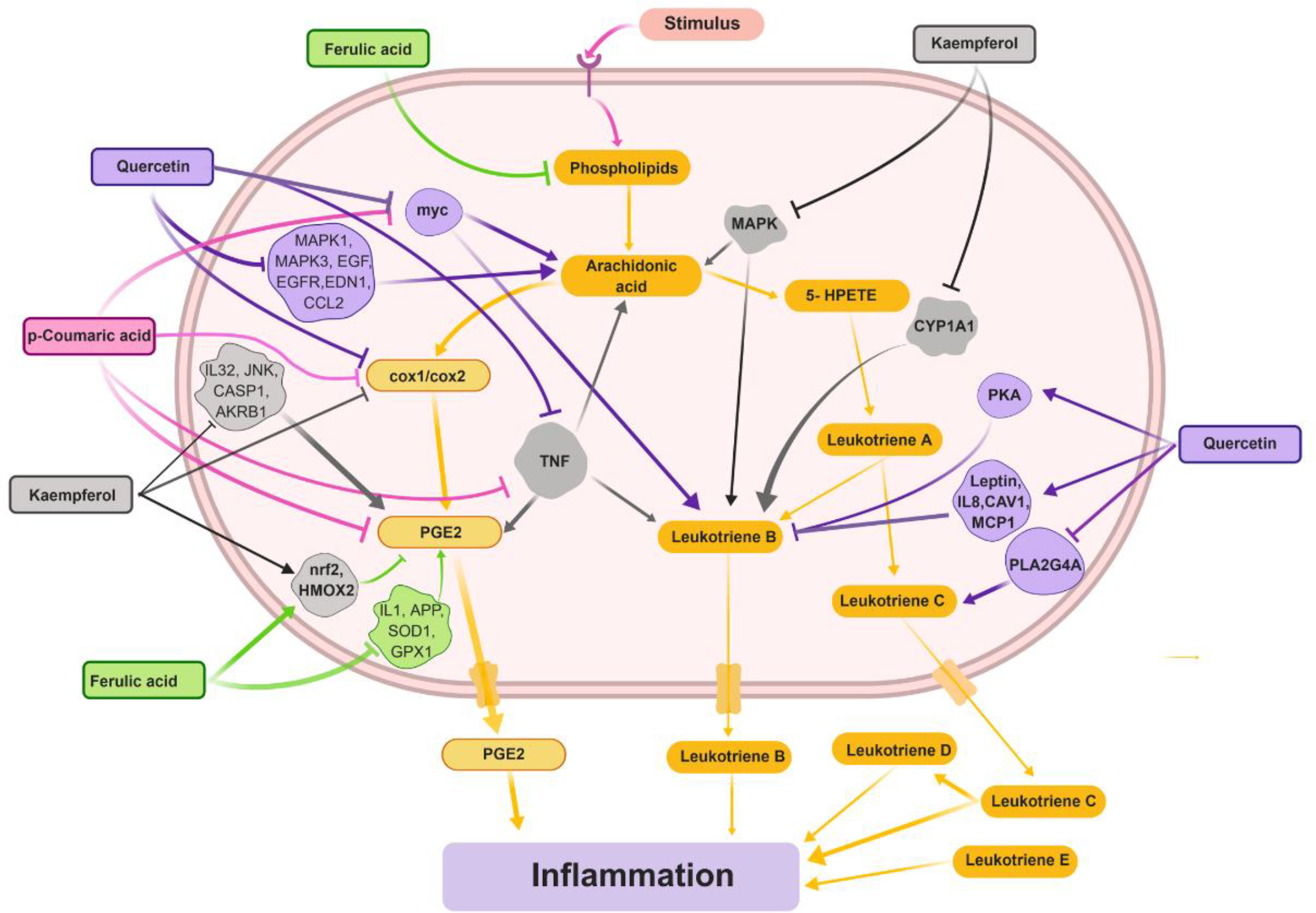
Possible mechanism of action of BC extracts to reduce inflammation in humans. Phenols and polyphenols, e.g., ferulic acid, p-coumaric acid, kaempferol, and quercetin in the BC extracts, inhibit the signaling molecules, stimulation of which may directly or indirectly result in inflammation. Proteins linked to inflammation participate in several canonical pathways underlying a range of biological activities. Ingenuity pathway analysis (IPA) identified that the BC extracts’ components could target many of these proteins (Table S1) and control inflammation. Arrows (*↓*) indicate direct stimulatory modification, whereas light up signs (┬) indicate an inhibitory modification. The Names of the proteins followed the Hugo nomenclature, NCBI Entrez, UniProtKB/Swiss-Prot system.

Ferulic acid, another component in the BC extract, negatively induces the synthesis of phospholipids. Phospholipid biosynthesis is critically important for the synthesis of arachidonic acids. Also, ferulic acids upregulate the expression of HMOX and nrf2, resulting in the reduction of prostaglandin E2 (PGE2). Moreover, all the compounds of the *Nigella* extract directly reduce the expression of cox1 and cox2, two essential factors for the synthesis of PGE2. Collectively, ferulic acid, quercetin, p-coumaric acid, and kaempferol help reduce inflammation by interfering with the extended arachidonic acid pathway (Figure 3; S1 Table).

## Discussion

In this study, we demonstrate the *in vitro* dose-response of the DMSO extract of BC to evaluate the anti-inflammatory and analgesic activities in the experimental models. The inhibition of the antigen-induced degranulation at the same concentration of 41 food samples was assayed in the rat basophilic leukemia RBL-2H3 cell line (data not shown). A further dose-response (40, 20, 10 and 5µg/mL) assay was carried out to confirm their anti-allergic activity of the bioactive compound as present in food samples and only DMSO, as well as AWA extract of BC, exhibited dose-response which support the concept that anti-allergic activity belongs to the phenolic compounds. Recent research findings showed that polyphenols, widely distributed among fruits, vegetables, and herbs, are widespread for their antioxidant capacity [29]. They also reported that some of them could exert anti-allergic activities [30] [31] [29].

Carrageenan-induced paw edema in rat models is increasingly being employed to determine anti-inflammatory effects in animal models. In response to carrageenan injection, the formation of edema occurs in a biphasic process. Mast cells around the damaged tissue released prostaglandin with histamine and serotonin mediate the first phase (1 - 2 h) of the carrageenan model. Bradykinin, leukotrienes, polymorphonuclear cells, and continuous secretion of prostaglandins from macrophages tissue mediate the second phage (3 - 5h) of the carrageenan model [32] [25]. Our study indicates that the DMSO extracts of BC exhibited significant (p < 0.05) inhibition of paw edema in rats at the doses of 200, 400, and 800 mg/kg, p.o in the second phases of inflammatory response. DMSO extracts of BC at the dose of 800 mg/kg produced the maximum inhibition of carrageenan-induced paw edema volume. The result of the present study showed the maximum inhibition of carrageenan-induced paw edema volume in rats by BC in the second phase of post carrageenan injection at 3-5 h, maybe due to the modulatory principles acting with the prostaglandin alley. Moreover, the present study supports the previous findings that BC seed polyphenol plays a crucial role as protective factors against

“Acetic acid-induced writhing experiment” is a well-known protocol for assessing the analgesic potency of medicinal products [33]. Intraperitoneal injection of acetic acid that triggers pain sensation is due to the prostaglandins and lipoxygenase products from arachidonic acid liberated from phospholipids by cyclooxygenase [34]. Thus, the significant (p<0.05) reduction of writhing in this study by DMSO extracts of BC at the doses of 200, 400, and 800 mg/kg, p.o suggest analgesic activity peripherally mediated through inhibition of prostaglandins and other endogenous pain mediators. The highest concentration (800 mg/kg) of the test samples exhibited a peak of analgesic effect significantly (p<0.05) in the acetic acid-induced writhing test. These results indicate the peripheral analgesic potential of the DMSO extracts of the test samples which could be exhibited due to the suppression of peritoneal surface receptors through inhibited cyclooxygenase activity. The current study findings show concordance with previous findings that BC seed polyphenol inhibits acetic acid-induced writhing in the mouse model [35].

Through an examination of the literature, we also note how to present findings confirm previous work in the field. Specifically, the *in vitro* and *in vivo* experiments performed in this study are in good agreement with those of prior experiments on BC’s anti-inflammatory activity, lending credence to the idea that BC is a potent medicinal plant for therapeutic uses.

In addition to the wet-lab analysis, ingenuity pathway analysis provides evidence for the role of the *Nigella* extracts in the reduction of inflammation. Earlier studies revealed that some molecules including myc, MAPK, EGFR, TNF, are involved in the cellular responses for inducing the inflammation [36] [37] [38] [39]. We have found that compounds in the *Nigella* extracts, including ferulic acid, quercetin, p-coumaric acid, and kaempferol, downregulates the inflammation-inducing signaling molecules in humans (Figure 3). The potential role of the activation of pKA, HMOX, and nrf2 pathways led to anti-inflammatory effects has been reported in many research groups [40] [41] [42] [43]. Our analysis uncovered that compounds in the *Nigella* extracts upregulate the pKA, HMOX, and nrf2 (Figure 3).

Furthermore, we have shown the potential aspects of the anti-inflammatory mechanism of BC that were previously only speculated upon. For example, Nasuti *et al*. found BC to reduce only the acute inflammation suggested that BC’s mechanism of action is tied to interleukins, TNF, and prostaglandins [44]. In the ingenuity pathway analysis, we have shown various components of BC act through IL32, IL8, IL1, TNF, and PGE2 to reduce inflammation. In another study, researchers discovered that BC could decrease either IL-6 levels of IL-1B levels, depending on its storage conditions [45]. These findings establish a connection between BC and interleukins in BC’s anti-inflammatory activity, which we have at least partially outlined in Figure 3. Similarly, other researchers speculated that BC WSE inhibits COX2 and that BC inhibits prostaglandin synthesis through interaction with COX-1 and COX-2 [46]. Both hypotheses are supported by the results of our Ingenuity pathway analysis, as depicted in Figure 3. Furthermore, Babar et al. suggested that phenols/polyphenols in BC might be responsible for its anti-inflammatory activity [46]. Through our pathway analysis, we have found that these phenols and polyphenols are ferulic acid, p-coumaric acid, kaempferol, and quercetin. It is speculated that the inhibitory effect on the secretion of leukotrienes and prostaglandins by thymoquinone may be responsible for BC’s anti-inflammatory activity [47]. We have shown that this is the case, although, instead of thymoquinone, quercetin and ferulic acid were found to be the components of BC responsible for this aspect of its anti-inflammatory mechanism. In another study, *N. sativa* showed superior activity in a screening of *Nigella* species for *in vitro* inhibition of PGE2 production catalyzed by COX1 and COX2 [48]. This lends credence to the mechanistic schematic we have depicted in Figure 3; as is shown in that diagram, several components of BC act to inhibit COX1/COX2 activity, which is indeed necessary to produce PGE2. In yet another study, researchers discovered that levels of erk are decreased upon *N. sativa* treatment of damaged rat lungs [49]. The erk repression discussed here is supported by our mechanistic analysis, which has found that kaempferol, a component of *N. sativa*, inhibits ERK1/2 activity. As evidenced by these examples from the literature, the results we have found build on previous research to provide crucial insight into the medicinal properties of a common natural product.

Together with the *in vitro* and *in vivo* studies, the *in silico* analysis in this study provides strong evidence that the bioactive compounds in BC extract act as an anti-inflammatory agent by regulating several downstream and upstream pathways of inflammation.

## Conclusion

In summary, the findings of the present study support that a BC extract has anti-inflammatory, anti-allergic, and analgesic activity. This study, involving *in vitro* and *in vivo* systems, indicates the possibility that BC extract might help treat pathologies that involve chronic inflammation and pain. Also, *in silico* analyses suggested that the phenolic compounds in BC extracts may play a key role in inhibiting inflammatory reactions in human? Besides, through the Ingenuity pathway analysis, the present study has supported and elaborated on suggestions found in the literature regarding BC’s biochemical mechanism of action. Thus, in understanding how to present findings relate to prior work in the field, we have gained insight into the vitality of this research for future efforts to study the anti-inflammatory effects of *N. sativa*. Future studies should focus on elucidating the mechanistic pathways through which compounds in BC extracts exert such pharmacologic effects to prevent inflammation.

## Supplementary Materials

Table S1: Components of the reactome *Nigella sativa* works on to impact on human inflammations.

## Acknowledgments

We thank Aftab Uz Zaman Noor, MSc (Université Claude Bernard Lyon 1, France) for his in-kind assistance for conducting the silico study.

## Author Contributions

Conceptualization, N.S. M.K.H., and S.A.; methodology, N.S., A.A., A.G., S.A., M.K.H. and M.S.A.P.; software, S.A., and M.S.A.P; validation and scrutinization, N.S., M.K.H., and U.P; investigation, N.S., A.A., A.G., S.A. and M.S.A.P.; formal analysis, N.S., A.A., A.G., S.A., and M.S.A.P.; writing—original draft preparation, A.A., A.G., and S.A.; writing—review and editing, N.S., M.K.H., and U.P.; supervision, N.S.; project administration, N.S., and M.K.H.; funding acquisition, N.S.

## Approval for animal experiments

Animals were housed at the Animal Housing Facility of Institute of Nutrition and Food Sciences, University of Dhaka, Bangladesh, following guidelines by their Ethics Committee for the Treatment of Laboratory Small Animals. All procedures were reviewed and approved by the Ethical and Animal Care Committee of the Biological Science faculty, University of Dhaka, Bangladesh. The committee’s reference number is not available.

## Approval for human experiments

Not applicable.

## Approval for experimental procedures

All experimental procedures were approved by the internal committee of Biological Science faculty headed by Dean of Bioscience faculty, University of Dhaka, Bangladesh.

## Informed consent

Informed consent was obtained from all individual participants included in the study.

## Data availability

Authors always welcome to share the data required for reviewers and other researchers.

## Abbreviations

BC: Black cumin
RBL: Rat basophilic leukemia
IPA: Ingenuity Pathway Analysis
AWA: Acetone-water-acetic acid
DMSO: Dimethyl sulfoxide

